# Identification and characterization of cis-regulatory elements for photoreceptor type-specific transcription in zebrafish

**DOI:** 10.1101/683284

**Authors:** Wei Fang, Yi Wen, Xiangyun Wei

## Abstract

Tissue-specific or cell type-specific transcription of protein-coding genes is controlled by both *t*rans-*r*egulatory *e*lements (TREs) and *c*is-*r*egulatory *e*lements (CREs). However, it is challenging to identify TREs and CREs, which are unknown for most genes. Here, we describe a protocol for identifying two types of transcription-activating CREs—core promoters and enhancers—of zebrafish photoreceptor type-specific genes. This protocol is composed of three phases: bioinformatic prediction, experimental validation, and characterization of the CREs. To better illustrate the principles and logic of this protocol, we exemplify it with the discovery of the core promoter and enhancer of the *mpp5b* apical polarity gene (also known as *ponli*), whose red, green, and blue (RGB) cone-specific transcription requires its enhancer, a member of the rainbow enhancer family. While exemplified with an RGB cone-specific gene, this protocol is general and can be used to identify the core promoters and enhancers of other protein-coding genes.

## 1. Introduction

In eukaryotes, tissue-specific or cell type-specific transcription of each protein-coding gene is regulated by specific *t*rans-*r*egulatory *e*lements (TREs) and *c*is-*r*egulatory *e*lements (CREs). Both TREs and CREs can be divided into multiple categories. TREs encompass RNA polymerase II, transcription factors, transcriptional coregulators, chromatin modifiers, regulatory RNAs, etc. (Latchman, 2008; Hughes, 2011; Latchman, 2015). And CREs encompass core promoters, enhancers, locus control regions, silencers, and insulators (Juven-Gershon and Kadonaga, 2010; Yanez-Cuna et al., 2013; Nelson and Wardle, 2013; Long et al., 2016). TREs and CREs regulate transcription through complex interactions. At the center of these interactions are the specific bindings between transcription factors and CREs (Juven-Gershon and Kadonaga, 2010; Long et al., 2016). Thus, identifying CREs is an essential step toward understanding the spatiotemporal transcription of a gene.

Each category of CREs plays different roles in transcription. Core promoters mediate the docking of the transcription pre-initiation complex by recruiting basic transcription factors; thus, they are required for transcription initiation. Core promoters are about 100–200 bp long and contain transcription start sites (TSSs) (Lenhard et al., 2012). By contrast, enhancers, spanning about 80–250 bp, recruit additional transcription factors to activate core-promoter-bound transcription initiation complex; thus, enhancers are required for transcription completion, often activating transcription in a spatiotemporally-specific manner (Kulaeva et al., 2012; Haberle and Stark, 2018). Unlike core promoters, enhancers locate away from TSSs, often thousands of base pairs away. However, in some genes, transcriptional specificity-deciding cis elements locate very close to the TSSs, around 100–200 bp in distance; these elements are sometimes referred to as promoter-proximal elements or proximal promoters in the literature (Lodish et al., 2000; Lenhard et al., 2012). Despite their proximity to TSSs, promoter-proximal elements or proximal promoters are functionally like enhancers (Lodish et al., 2000). For the sake of simplicity and clarity, here we encompass promoter-proximal elements or proximal promoters in the category of enhancers per their function of stimulating transcription. Thus, it needs to be kept in mind that the compositions and locations of enhancers can vary drastically from gene to gene.

Like enhancers, locus control regions activate specific transcription, but they exert broader effects by controlling a cluster of related genes within a locus (Bulger and Groudine, 1999). In contrast to enhancers, silencers suppress transcription by recruiting transcription suppressors (Chong et al., 1995; Nelson and Wardle, 2013). Unlike enhancers, locus control regions, and silencers, all of which act directly on their target genes, insulators act indirectly by preventing enhancers, silencers, and locus control regions on one side of an insulator from affecting genes on the opposite side of the insulator (Herold et al., 2012). These CRE properties suggest that core promoters and enhancers are the basic transcription-activating elements that mediate spatiotemporal transcription of a gene, making these two CREs useful for practical applications, such as expressing therapeutic genes in specific cell types in gene therapies. Despite their importance, the core promoters and enhancers of most genes remain unknown and challenging to identify, particularly enhancers.

Here, we describe a protocol for identifying the core promoters and enhancers of zebrafish photoreceptor type-specific genes. This protocol is composed of three phases: Phase I predicts the CREs with bioinformatic algorithms; Phase II validates the candidate CREs in vivo with transgenic expression assays; and Phase III characterizes the function of CREs and their motifs by element-swapping and sequence motif-mutating analyses. To better illustrate the principles and the logic of this protocol, we exemplify the protocol with the discovery of the core promoter and enhancer of the *mpp5b* zebrafish apical polarity gene (also known as *ponli*; Zou et al., 2010). *mpp5b* is restrictively expressed in red, green, and blue (RGB) cones, and this specific transcription needs its enhancer, which is a member of the rainbow enhancer family (Fang et al., 2017). Although exemplified with an RGB cone-specific gene, this protocol has broad applications and can be used to identify the core promoters and enhancers of other tissue-specific or cell type-specific protein-coding genes.

## 2. Materials

### 2.1 Fish lines and fish care

Zebrafish strains can be purchased from Zebrafish International Resource Center (https://zebrafish.org/home/guide.php). Medaka fish can be obtained from Arizona Aquantic Garden (https://www.azgardens.com/). Zebrafish and medaka can be maintained on a 14-hr light/10-hr dark cycle. Please follow institutional regulations on experimental animal care.

### 2.2 Bioinformatics tools

2.2.1 The UCSC Genome Browser (http://genome.ucsc.edu/) can be used to browse, extract, and compare genomic sequences of many model species.

2.2.2 The MEME Suite (http://meme-suite.org/), a bioinformatics suite of motif-based sequence analysis tools, can be used to identify short nucleotide or peptide sequence motifs and to search for them in long DNA or amino acid sequences.

2.2.3 TRANSFAC Professional (http://gene-regulation.com/), a web-based database of the known transcription factors, can be used to predict binding sites for known transcription factors in DNA sequences.

### 2.3 Reagents for recombinant DNA technologies

2.3.1 The Mini-Genomic DNA Buffer Set (Qiagen, Cat #: 19060) can be used to isolate the genomic DNAs from animal tissue samples.

2.3.2 The Platinum PCR SuperMix High Fidelity DNA polymerase (Invitrogen, Cat #: 12532016) can be used to amplify DNA fragments with a high degree of accuracy.

2.3.3 The QIAGEN Plasmid Mega Kit (Cat #: 12181) can be used to isolate DNA from BAC or PAC clones.

2.3.4 The Q5 Site-Directed Mutagenesis Kit (NEB, Cat #: E0554S) can be used to make deletion and substitution mutations of DNA constructs.

2.3.5 I-SceI, a homing endonuclease that recognizes and cuts the target sequence of TAGGGATAACAGGGTAAT (NEB, Cat#: R0694S), is recommended to be coinjected with the pSceI-based transgenic constructs to generate transgenic zebrafish (Thermes et al., 2002).

## 3. Equipment

3.1 A Flaming/Brown Micropipette Puller (Model P-97; Sutter Instrument Co.) can be used to prepare embryo injection needles with borosilicate glass capillaries (Kwik-FilTM; World Precision Instruments, Inc. Cat #:1B100-4) (Yuan and Sun 2009; Rosen et al., 2009).

3.2 A Pneumatic PicoPump (World Precision Instruments, Inc.; PV820) can be used to inject DNA constructs into zebrafish embryos (Yuan and Sun 2009; Rosen et al., 2009).

## 4. Methods

### 4.1 Phase I: Bioinformatic prediction of the core promoter and enhancer of a zebrafish gene

#### Rationale

Core promoters and enhancers can be predicted with many bioinformatic algorithms. Such prediction is based on the logic that the DNA sequences that possess characteristic properties of CREs are likely CREs. The following three characteristic properties are particularly important for predicting core promoters and enhancers: locations, sequence conservation, and possession of transcription factor binding sites.

##### The locations of core promoters and enhancers

Core promoters contain TSSs (Juven-Gershon and Kadonaga, 2010; Lenhard et al., 2012;Hardison and Tylor, 2012; Haberle and Stark, 2018). By contrast, enhancers do not overlap with TSSs and often reside thousands of base pairs away from TSSs, up to 100 kb in distance. (In some genes, enhancers may also reside not far from TSSs, like 100–200 bp away from TSSs.) Enhancers can localize either upstream or downstream of the TSSs (Kulaeva et al., 2012). When residing downstream of a TSS, an enhancer can either locate within an intron or downstream of the gene. This versatility of enhancer locations makes them more challenging to identify than core promoters.

##### Sequence conservation of core promoters and enhancers

Core promoters and enhancers generally reside in conserved noncoding regions, thus flanked by un-conserved DNA sequences. Such sequence conservations exist both among orthologs of multiple species and among coregulated different genes within a single genome (Hardison, 2000; Pennacchio and Rubin, 2001; Elgar, 2009; Vavouri and Lehner, 2009).

##### Possession of binding sites for transcription factors

Core promoters and enhancers are densely-packed with highly conserved short sequence motifs (between 6–17 bp), some of which are palindromic or direct repeats. These sequence motifs are binding sites for transcription factors (Juven-Gershon et al., 2010; Goodrich and Tjian), which often bind to DNA as dimers or even trimers, with each transcription factor binding to a sequence as short as 3 nucleotide residues (Panne et al., 2007; Latchman, 2008; Hughes, 2011). For example, an AT-rich TATA box, normally residing 30 bp upstream of the TSSs of some core promoters, recruits TBP of the TFIID complex (Nikolov et al., 1992), and an initiator, spanning the TSSs, recruits a TAFII250– TAFII150 dimer (Chalkley et al., 1999). Similarly, multiple binding sites in enhancers recruit various transcription factors which cooperate to mediate tissue-specific or cell type-specific transcription (Panne et al., 2007).

The above-mentioned properties can be used to predict core promoters and enhancers with bioinformatic tools via one or both of two strategies: The first strategy utilizes sequence conservation among the CREs of coregulated but different genes in a single genome (Beer and Tavazoie, 2004; Middendorf et al., 2004; Yuan et al., 2007; Warner et al., 2008; Rouault et al., 2014). While powerful, this strategy requires prior knowledge of co-transcription profiling, which might be missing or incomplete for the genes of interest. In addition, this strategy works less successfully in higher organisms than in yeast and other lower organisms, presumably because the CREs in higher organisms are often not restricted to nearby upstream sequences and because their motif organizations are more complex.

The second strategy, namely the phylogenetic footprinting strategy (Tagle et al., 1988), utilizes sequence conservation among the CREs of the orthologous genes of multiple species (Das and Dai, 2007). Although independent of prior knowledge of co-transcription profiling as required for the first strategy, the phylogenetic footprinting strategy can also run into problems when the orthologous sequences are either too closely-related, which makes a global multiple alignment uninformative, or too distantly-related, which could make it impossible to align conserved motifs (Das and Dai, 2007). However, with the genomic sequences of more species becoming available, the phylogenetic footprinting strategy becomes more and more practical.

Thus, Phase I of this protocol takes the phylogenetic footprinting strategy to predict the core promoter and enhancer of a zebrafish gene with web-based algorithms (Fig. 1). In the following, we exemplify the procedures of such prediction with the *mpp5b* gene.

**Figure 1.**
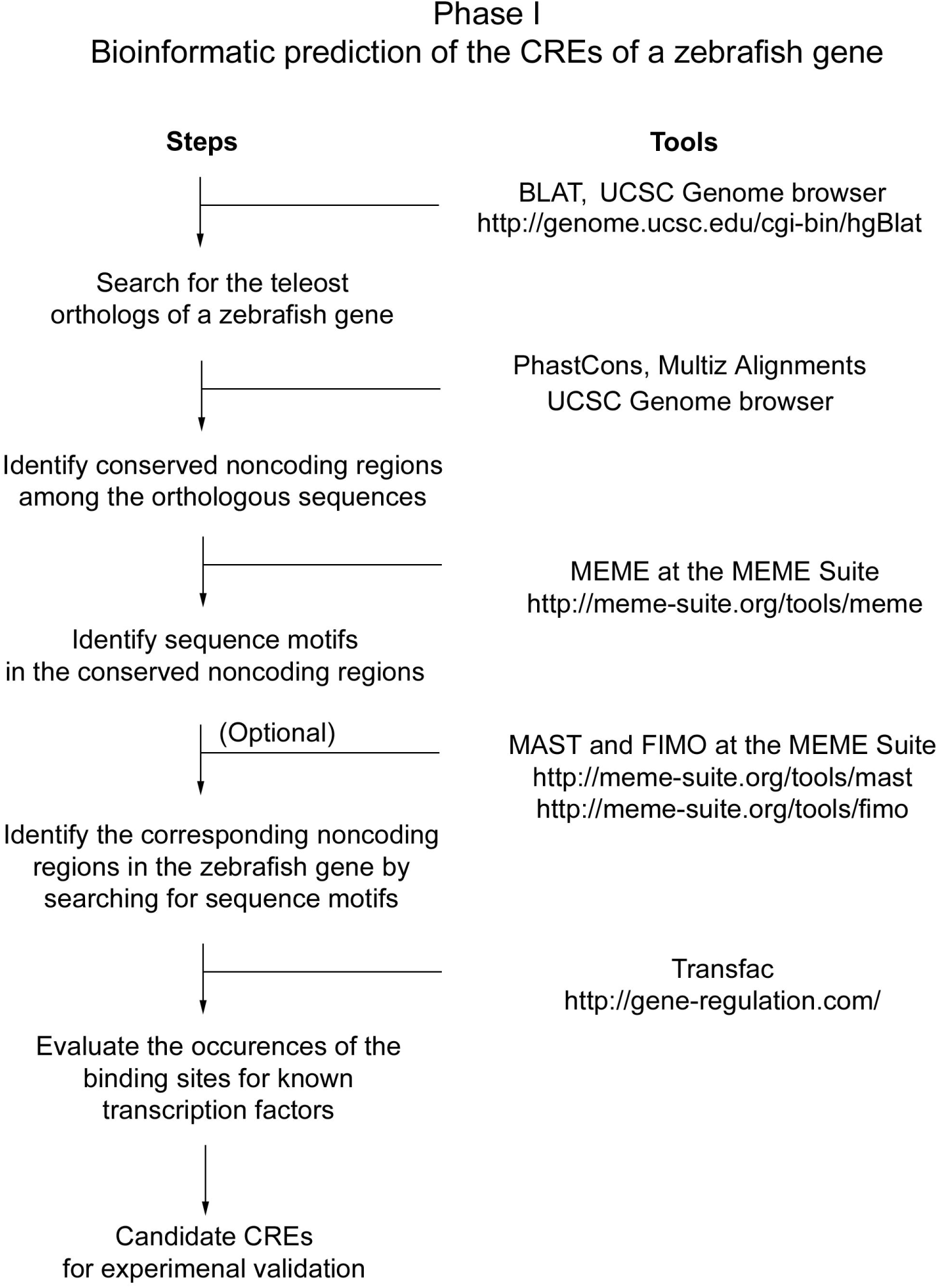
The outline of Phase I of the protocol—CRE prediction. The flow chart shows the steps to predict CREs with some web-based bioinformatic tools.

#### Procedures

##### 4.1.1 Identify teleost orthologs of a zebrafish gene

Several teleost orthologs of a zebrafish gene can be identified at the UCSC genome browser with the BLAT algorithm, which searches for genomic regions that are homologous to query sequences (http://genome.ucsc.edu/; Kent et al., 2002a; for an introduction to the UCSC genome browser, see Kent et al., 2002b; for video tutorials on the UCSC genome browser, visit http://genome.ucsc.edu/training/index.html).

For example, to locate the teleost orthologs of the zebrafish *mpp5b* gene, open the BLAT search page at http://genome.ucsc.edu/cgi-bin/hgBlat, which can also be reached by clicking on the “BLAT” button at the home page of the USCS genome browser. Then select “Medaka” from the genome options and leave other parameters in their default settings. (Searching the medaka genome works better for *mpp5b* than searching other fish genomes because the orthologs of more teleost species were aligned, thus maximally revealing sequence conservations. For other genes, we recommend trying all available non-zebrafish teleost genomes for the best outcomes. Please note that teleost genomes used for evolutionary conservation analyses might have not been updated.) Next, paste the entire amino acid sequence of the zebrafish Mpp5b protein (gene accession number GU197553) in the query sequence box (please note that amino acid sequences serve better than nucleotide sequences as queries for cross-species BLAT searches), then click on the “submit” button to reveal two medaka genes to be identified, one on chromosome 24 with higher matching scores than the other, on chromosome 20 (Fig 2A). Then click on the “browser” button of the gene with higher scores (i.e., the medaka ortholog of zebrafish *mpp5b*) to display the medaka genomic region, which the *mpp5b* gene matches (Fig. 2B). (Please note that the amino acid query sequence of zebrafish Mpp5b is split into sections and matched only to the coding regions of the predicted medaka *mpp5b* gene. The noncoding sequences of the medaka *mpp5b* mRNA is not marked because medaka *mpp5b* has yet to be verified experimentally and to be annotated as such.)

**Figure 2.**
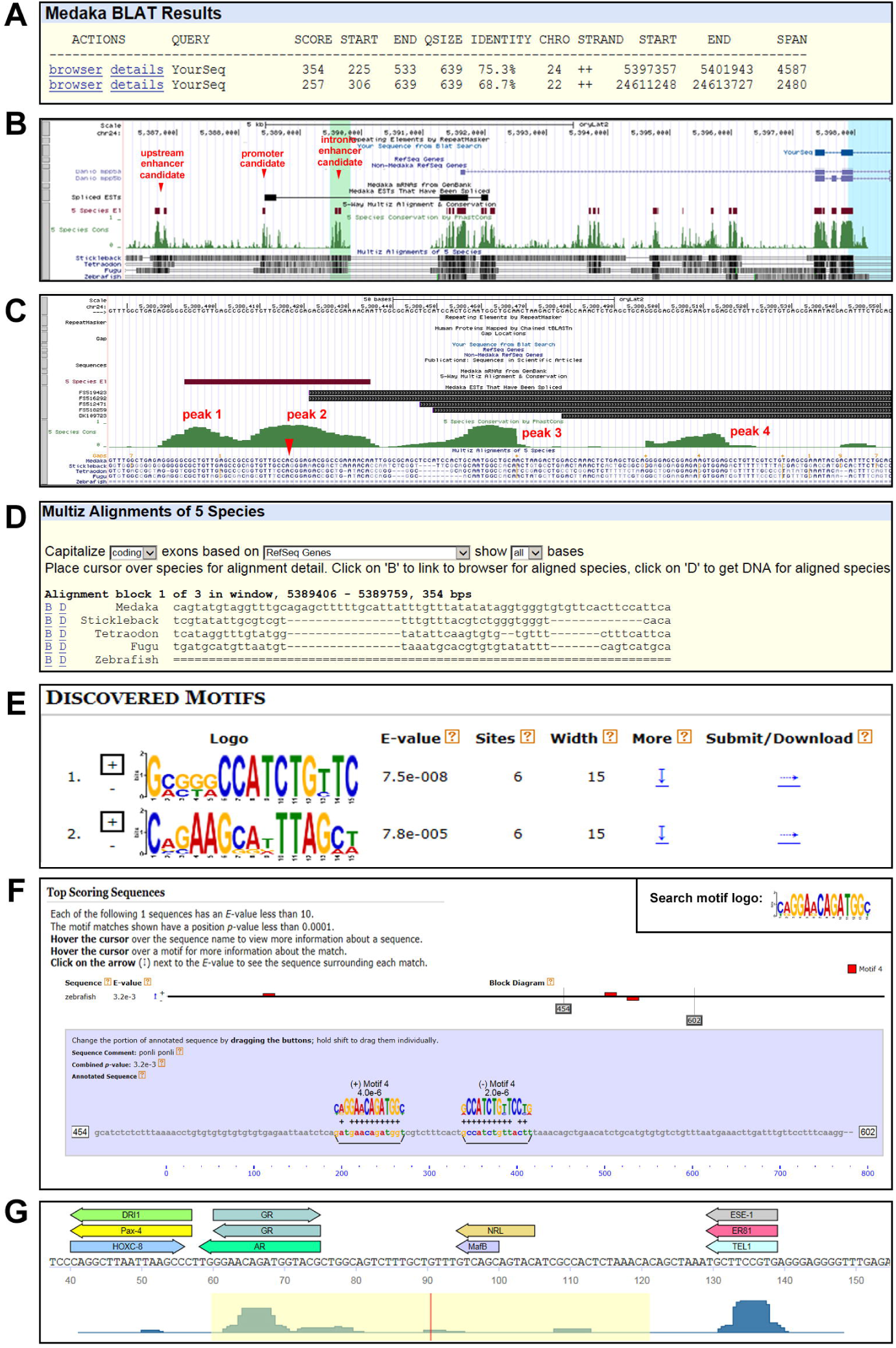
Example results of the bioinformatic analyses. **A**. A result page of a BLAT search of the medaka genome with the amino acid sequence of zebrafish Mpp5b protein as the query, showing two medaka homologs identified. **B**. A user interface page at the UCSC genome browser displays the query of zebrafish Mpp5b aligned to the medaka genome, EST clones of the medaka *mpp5b* ortholog, published zebrafish *mpp5a* and *mpp5b* genes aligned to the medaka genome, 5 Species Conservation by PhastCons, Multiz Alignments of 5 species, etc. (More tracks of annotations can be made available by adjusting the track settings.) Red arrowheads indicate the locations of some core promoter and enhancer candidates of the *mpp5b* orthologs. **C**. A page at the UCSC genome browser displays the region around the transcription start site (red arrowhead) of the *mpp5b* orthologs, showing the four conservation peaks by PhastCons (peaks 1–4). Hyphens, sequence gaps; equal signs, un-alignable regions. **D**. A page at the UCSC genome browser displays the Multiz Alignments of the *mpp5b* orthologs and the links to browse and download the sequences of a selected genomic region. **E**. A result page of an MEME analysis displays the consensus sequences of two of many motifs in the enhancers of *mpp5b* genes. **F**. A result page of a MAST analysis shows that three sequence motifs (red boxes) in a zebrafish sequence were found to be similar to the query motif (motif 4, inset). **G**. A result page of a MATCH analysis in Transfac shows that the tilapia *mpp5b* enhancer is predicted to contain binding sites for known transcription factors.

##### 4.1.2 Identify the TSS of the gene of interest

It is important to identify TSSs because they are the dividing points between the upstream and downstream sequences of genes. However, TSSs need to be extrapolated from the 5’ ends of mRNA sequences, which can be obtained experimentally by 5’RACE analyses (the 5’ RACE System for Rapid Amplification of cDNA Ends, version 2.0; Thermo Fisher Scientific, Cat #: 18374058). Alternatively, one can estimate TSSs from the 5’ end sequences of ESTs (expressed sequence tags). Please note that EST sequences may not extend to TSSs; thus, they may not pinpoint TSSs. Please also note that even if ESTs actually cover TSSs, different ESTs may suggest different TSSs because the core promoter of a gene can be a dispersed type of core promoter which possesses multiple close-spaced TSSs (Carninci et al., 2006; Lenhard et al., 2012).

For example, to estimate the TSS of the medaka *mpp5b* gene, whose full-length cDNA sequence has not been published, we can utilize its EST sequences. Two sets of EST sequences are available, one set for the 5’ end of the gene and the other for its 3’ end. Drag the interactive screen of the UCSC genome browser to the upstream direction to reveal the five 5’ end ESTs (FS519423, FS516292, FS12471, FS518259, and DK109723) (Fig. 2C). Among them, FS519423 and FS516292 extend farthest in the upstream direction, making their 5’ ends the best estimate of the TSS, although the TSS of zebrafish *mpp5b* (Zou et al., 2010) and the sequence conservation of the TSS may suggest the TSS of medaka *mpp5b* to be 3 and 4 bp upstream of the FS519423 and FS516292, respectively. Despite this uncertainty, the 5’ end ETS sequences provide a very close estimate of the TSS, thus revealing the region of the core promoter.

##### 4.1.3 Identify conserved noncoding regions among orthologous sequences

Once the TSS has been estimated or pinpointed, the conserved upstream and downstream noncoding regions are revealed by high values in the PhastCons histograms of the evolutionary conservation and Multiz alignments of the five fish species of medaka, tetraodon, fugu, stickleback, and zebrafish on the UCSC genome browser. (Please note that coding regions are also highly conserved, so the conserved noncoding regions need to be distinguished from the coding regions, which normally align with the query amino acid sequences.) The conserved noncoding regions within 100 bp upstream and downstream of TSSs likely contain the core promoter; by contrast, more distal conserved noncoding regions, particularly in a single stretch of 80–250 bp, whether upstream or downstream of the TSS, likely harbor enhancers.

For example, the PhastCons Conservation histograms and Multiz alignments revealed that within 150 bp of the TSSs of teleost *mpp5b* genes (from −45 to +105), four short sequence motifs are conserved among medaka, stickleback, tetraodon, and fugu (please note that zebrafish sequence could not be aligned with other sequences at the TSS region, likely due to divergence in sequence and motif configuration) (Fig. 2C). Together, these four short motifs likely constitute the core promoter of the *mpp5b* genes. In addition, PhastCons Conservation and Multiz alignments also revealed conserved noncoding regions in the upstream intergenic and downstream intronic regions, which are candidates for the enhancer of *mpp5b* (Fig. 2B for two examples).

Once conserved noncoding regions are identified, their sequences can be downloaded by step 4.1.4, their sequence motifs can be analyzed by step 4.1.5, and the possession of binding sites for known transcription factors can be assessed by step 4.1.7. But if the corresponding conserved noncoding regions of the zebrafish gene are not revealed by PhastCons Conservation and Multiz alignments, as in the case of *mpp5b*, follow step 4.1.6 to identify these regions.

##### 4.1.4 Download the sequences of the conserved noncoding regions

First select the regions in the interactive page to zoom into the conserved noncoding regions, then click on the black bars in the information track of “Multzi alignment of 5 species” (Fig 2B, at the bottom) to open sequence alignments as well as the links for downloading the sequences in the FASTA format (Fig. 2D, showing a section of the *mpp5b* intron 1).

##### 4.1.5 Identify conserved sequence motifs

To search the conserved noncoding regions for sequence motifs, we recommend using the MEME algorithm (a component of the MEME suite; http://meme-suite.org/; Bailey et al., 1994; Bailey et al., 2009). Open the MEME website at http://meme-suite.org/tools/meme, input the sequences of the orthologous conserved noncoding regions in the FASTA format, then in the advanced options, choose the ranges of motif lengths arbitrarily between 6 and 17 to perform a search. We recommend searching repeatedly with different motif lengths because transcription factors bind to short motifs of different lengths and MEME does not detect motifs outside the selected length ranges. Besides identifying conserved motifs, MEME also determines the consensus sequences of the identified motifs, which can be downloaded in the “Minimal MEME” format (Fig. 2E, clicking on the horizontal arrows to download) for later MAST and FIMO analyses as described below.

4.1.6 Identify corresponding noncoding regions in a zebrafish gene: To determine if a zebrafish DNA sequence contains regions that correspond to the conserved noncoding regions identified in other teleost fish by PhastCons Conservation and Multiz alignments, one can use the MAST and FIMO algorithms to search a candiate DNA sequence for short conserved sequence motifs identified by MEME. MAST and FIMO algorithms are designed to determine the occurrence and location of query motifs in DNA sequences (components of MEME suite, http://meme-suite.org/) (Bailey and Gribskov, 1998; Grant et al., 2011). If a small region in the DNA sequence (e.g., 80–250 bp) is clustered with multiple short motifs identified previously by MEME, this region is likely the equivalent conserved noncoding region in zebrafish.

For example, to determine if the first intron of the zebrafish *mpp5b* gene contains a region that corresponds to the candidate intronic enhancer identified in other fish species by PhastCons Conservation and Multiz alignments at step 4.1.3 (Fig. 2B, right red arrowhead), open the MAST page at http://meme-suite.org/tools/mast, enter the sequence of the first intron of the zebrafish *mpp5b* gene (Genomic sequences can be downloaded at the UCSC genome browser by clicking on the “View” tab and then the “DNA” link) and then individual motifs identified by MEME in the “Minimal MEME” format to perform the analysis. Figure 2F reveals the example results of searching a region within the first intron of zebrafish *mpp5b* for a single motif, noting that three occurrences of the motif were identified (red boxes). To increase the chance of identifying a conserved noncoding region, we recommend first searching with highly conserved motifs to narrow down the target regions and then searching with less conserved motifs within the narrower regions. This motif-searching analysis can also be performed with the FIMO algorithm (http://meme-suite.org/tools/fimo). Ultimately, an intronic region in the middle of the first intron of the zebrafish *mpp5b* gene was identified to contain multiple conserved sequence motifs which were identified in other fish species by MEME, and this region was eventually confirmed to harbor the enhancer of zebrafish *mpp5b* (Fang et al., 2017).

##### 4.1.7 Evaluate the presence of binding sites for known transcription factors in conserved noncoding regions

Authentic CREs contain binding sites for transcription factors. Thus, the presence of binding sites for known transcription factors in conserved noncoding regions would strongly suggest that they are authentic CREs. To evaluate whether conserved noncoding regions contain potential transcription factor binding sites, one can use TRANSFAC Professional (http://gene-regulation.com/), whose database contains experimentally-verified information on eukaryotic transcription factors. (An institutional registration of TRANSFAC is required to access their most updated database.)

For example, to search for potential transcription factor binding sites in the intronic enhancer candidate of the medaka *mpp5b* gene, open “MATCH tool” on the TRANSFAC webpage, then enter the conserved intronic sequence as an input and click on “start search” to reveal a cluster of motifs that are potential bindings sites for some transcription factors (Fig. 2G). (Please note that whether or not these transcription factors actually bind to the intronic sequence needs to be determined by experiments.)

Through the above steps, candidate core promoters and enhancers are predicted in Phase I. Please keep in mind that prediction is not 100% accurate. Therefore, the predicted candidate core promoters and enhancers need to be experimentally validated in Phase II of this protocol.

### 4.2 Phase II: Experimental validation of core promoters and enhancers

#### Rationale

In Phase II, candidate core promoters and enhancers need to be validated experimentally for three reasons: (1) The bioinformatic algorithms used for CRE prediction are not perfect, and making things worse, these imperfect algorithms are based on our incomplete understanding of transcriptional regulation; (2) conserved noncoding sequences do not always carry transcriptional activities; (3) functionally conserved CREs do not always display conservation in sequence homology because transcription factors can bind to degenerative sequences and because the number, location, and orientation of transcription factor binding sites can be flexible, thus evading homology detection (Wittkopp and Kalay, 2011; Nelson and Wardle, 2013). The transcriptional activities of candidate core promoters and enhancers can be assessed in transgenic zebrafish in vivo by three steps (Fig. 3A): First, generate transgenic constructs that use candidate CREs to control the expression of a fluorescent protein reporter gene, such as a GFP (green fluorescent protein) gene; second, inject the constructs into zebrafish embryos and raise the injected embryos to desired developmental stages; third, examine the expression patterns of the reporter gene in these transgenic fish to assess the transcriptional activities of the candidate CREs.

**Figure 3.**
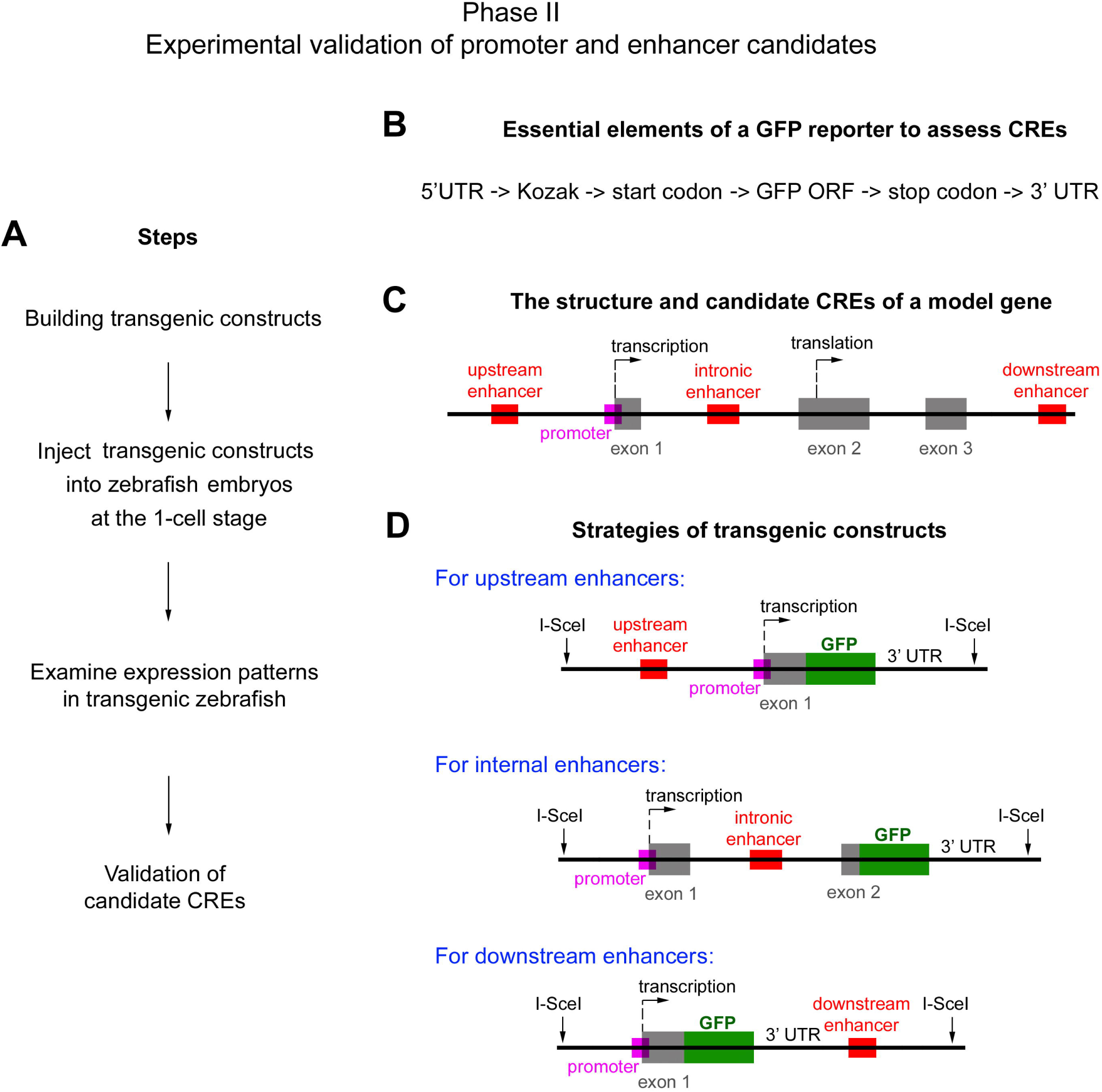
The outline of Phase II of the protocol—experimental validation of core promoter and enhancer candidates. **A**. The flow chart illustrates the steps to validate candidate core promoters and enhancers. **B**. Some of the essential elements of a GFP reporter gene are presented from left to right in the upstream-to-downstream order in which they shall reside in the reporter gene. **C**. The diagram illustrates the locations of the core promoter (a magenta box) and three possible locations of the enhancer of a model gene (red boxes, upstream, intronic, and downstream positions). Arrows indicate the transcription and translation start sites. Grey boxes stand for the exons of the model gene. **D**. Diagrams illustrate three strategies for making transgene constructs to accommodate the three possible locations of enhancers.

Phase II of the protocol utilizes a variety of techniques of molecular and cellular biology, such as PCR, restriction digestion, DNA ligation, plasmid isolation, plasmid construction, *E. coli* transformation, immunohistochemistry, microscopy, etc. For detailed explanations of these basic techniques, please refer to “*Molecular Cloning: A Laboratory Manual*” (Green and Sambrook, 2012) and “Immunohistochemistry: Basics and Methods” (Buchwalow and Bocker, 2010). Here, we explain only the outline of Phase II and some precautions pertinent to the purpose of this protocol.

#### Procedures

##### 4.2.1 The general elements of a transgenic reporter construct

For a GFP reporter gene to be specifically expressed, the transgene construct must have several essential elements besides a core promoter and an enhancer. These elements should be arranged from the upstream end to the downstream end in the following order: 5’ untranslated region (5’ UTR), Kozak sequence, start codon, the open reading frame (ORF) of GFP, stop codon, 3’ untranslated region (3’ UTR, containing a polyadenylation signal) (Fig. 3B). Many GFP expression vectors have these elements, so they can be modified to make a transgene reporter construct. For the basic properties of these elements, please refer to textbooks “*Levin’s Gene XII*” and “*Gene Control*” (Krebs et al., 2017; Latchman, 2015). Besides maintaining the basic properties of these elements, the following specific precautions can also be used:

###### 5’ UTR

We recommend using the 5’ UTR of the gene of interest as the 5’ UTR of the transgenic reporter gene because 5’ UTRs likely contain core promoter motifs. Of course, when performing CRE-swapping experiments (see Phase III), 5’ UTRs of other genes can be used to create various combinations of core promoters and enhancers to assess their transcriptional activities.

###### Kozak sequence

To ensure effective translation, a Kozak sequence (consensus sequence: GCCA/GCCATGG) needs to be included before the start codon. However, if the entire 5’ UTR of the gene of interest is to be included in a transgene construct, it is not necessary to include a synthetic Kozak sequence even if a Kozak consensus sequence is not apparent in the 5’ UTR because a functional substitute of the Kozak sequence should exist in the 5’ UTR.

###### 3’ UTR

The SV40 3’ UTR can be used as the 3’ UTR for a reporter transgene because many GFP expression vectors use the SV40 3’ UTR, which contains a polyadenylation signal. Of course, the 3’ UTR of the gene of interest can also be used.

###### I-SceI sites

We recommend the I-SceI meganuclease-based transgenesis (Thermes et al., 2002). Thus, an I-SceI site needs to be placed at both the upstream and downstream boundaries of the transgene cassette (Fig. 3D).

##### 4.2.2 Integrate candidate core promoters and enhancers into transgene constructs

Like the above-mentioned essential elements of a gene, core promoters and enhancers also have their specific positions in a gene. The core promoter is positioned at the 5’ end of the gene. To ensure the entire core promoter is included in the construct, include some additional upstream sequence, 100–200 bp or thereabouts, along with conserved core promoter elements. (This precautious measure implies the possibility of including those enhancers that reside very close to TSSs in the core promoter candidates, thus making it more necessary to characterize CREs in the Phase III of the protocol to more accurately define the regions for the core promoter and enhancer.)

Unlike core promoters, the enhancer of a gene can have three possible locations: upstream, intronic, and downstream (Fig. 3C). When integrating an enhancer candidate into a transgene construct, we recommend preserving its natural position and orientation because it is yet unclear what roles the location and orientation of enhancers play in their transcriptional activity, even though conventional wisdom, which is based on a limited number of case studies, suggests that enhancers can regulate transcriptional activities regardless of location and orientation (Latchman, 2015). Accordingly, to minimize the risk of overlooking an enhancer, three types of constructs are devised to test candidate enhancers at the three different locations (Fig. 3D). Of course, after an enhancer has been validated, it would be interesting to assess how the alteration of its location and orientation affects its transcriptional activities.

When testing candidate enhancers, include some flanking non-conserved sequences as well so as to reduce the risk of missing less-conserved but functionally-important motifs. Subsequently, the enhancer region can be trimmed by making deletion mutations with the Q5 Site-Directed Mutagenesis Kit (NEB, Cat #: E0554S). For example, when identifying the intronic enhancer of *mpp5b*, all of intron 1 was first assessed. Then un-conserved sequences were removed to narrow down the enhancer region (Fang et al., 2017). When trimming an intronic enhancer, preserve the 5’ and 3’ splicing sites as well as the splicing branching site, which normally localizes 18–40 bases upstream of the 3’ splicing site (Latchman, 2015).

##### 4.2.3 Obtain the DNA fragments of candidate CREs

The sizes of candidate core promoters and enhancers are normally only a few hundred base pairs long. Thus, the DNA fragments of these candidate CREs can be amplified by PCR. High fidelity DNA polymerase, such as the Platinum PCR SuperMix High Fidelity DNA polymerase (Invitrogen, Cat #: 12532016), can be used for the PCR amplification. Template DNAs for PCR can be isolated from fish tissues with the Mini-Genomic DNA Buffer Set (Qiagen, Cat #: 19060). Alternatively, if large pieces of DNA fragments are to be tested, they can also be isolated from a BAC or PAC DNA clone by restriction digestion or PCR amplification. BAC and PAC clones are available from the BAC/PAC Resources Center (https://bacpacresources.org/). BAC and PAC DNA can be purified with a QIAGEN Plasmid Mega Kit (Cat #: 12181).

##### 4.2.4 Embryonic injection of transgene constructs

50 pg of a transgenic construct can be co-injected with 0.1 unit of I-SceI meganuclease (NEB, R0694S) into zebrafish embryos at the 1-cell stage. Detailed procedures of needle preparation and microinjection are illustrated by two video articles (Yuan and Sun, 2009; Rosen et al., 2009). After injection, raise the embryos in E3 medium (5 mM NaCl, 0.17 mM KCl, 0.33 mM CaCl_2_, 0.33 mM MgSO_4_, 0.00001 % (w/v) Methylene Blue) until 5 dpf (days postfertilization). To examine GFP expression in the eyes, melanin pigmentation needs to be blocked with 0.0003% phenylthiourea (PTU) in the E3 medium. When necessary, transfer the fish larva to an aquatic system to raise to desired developmental stages, such as 9 dpf or adulthood. The GFP expression patterns of the fish can be analyzed by confocal immunohistochemistry as follows.

##### 4.2.5 Evaluate GFP expression patterns by immunohistochemistry

Zebrafish retina has one type of rod photoreceptor and four types of cone photoreceptors (red, green, blue, and UV cones). The resulting GFP expression patterns in the photoreceptors of either the adult retina or the larval retina can be determined by confocal immunohistochemistry per the morphological characteristics and immunoreactivities as follows:

###### In the adult zebrafish retina, five criteria can be used to distinguish the types of photoreceptors that express GFP

(1) The regular planar patterning of zebrafish photoreceptors. In zebrafish, green, red, and blue cones coalesce into pentamers in the order of G-R-B-R-G within each row of photoreceptors, with a UV cone separating two cone pentamers (Robinson et al., 1993; Raymond et al., 1995; Zou et al., 2018); between neighboring rows, the positions of pentamers shift by three cells, generating alternating columns of blue-UV cones and columns of red-green cones. Unlike cones, the inner segments of rod photoreceptors cluster around UV cones, and the cross sections of their inner segments are very thin at the level of RGB cone nuclei (Fig. 4 A; Zou et al., 2012; Fang et al., 2013; Table 1); (2) The DAPI staining of the nuclei of RGB cones and UV cone is lighter than that of rod nuclei (Fig. 4B; Table 1); (3) The nuclei of different types of photoreceptor localize to distinct regions relative to the outer limiting membrane (OLM), with rod nuclei basal and distal to the OLM, UV cone nuclei basal to and juxtaposing the OLM, and elongated RGB cone nuclei apical to and juxtaposing the OLM (Fig. 4B; Table 1); (4) Immuno-reactivity to photoreceptor type-specific antibodies or (Tables 1, the opsin antibodies, Vihtelic et al., 1999; Zpr1 antibody for double cones—red and green cones, ZFIN; Zpr3 antibody for rods, ZFIN); (5) Comparison with existing transgenic zebrafish lines, in which various types of photoreceptors are highlighted by transgenic proteins (Table 2).

**Table 1.**
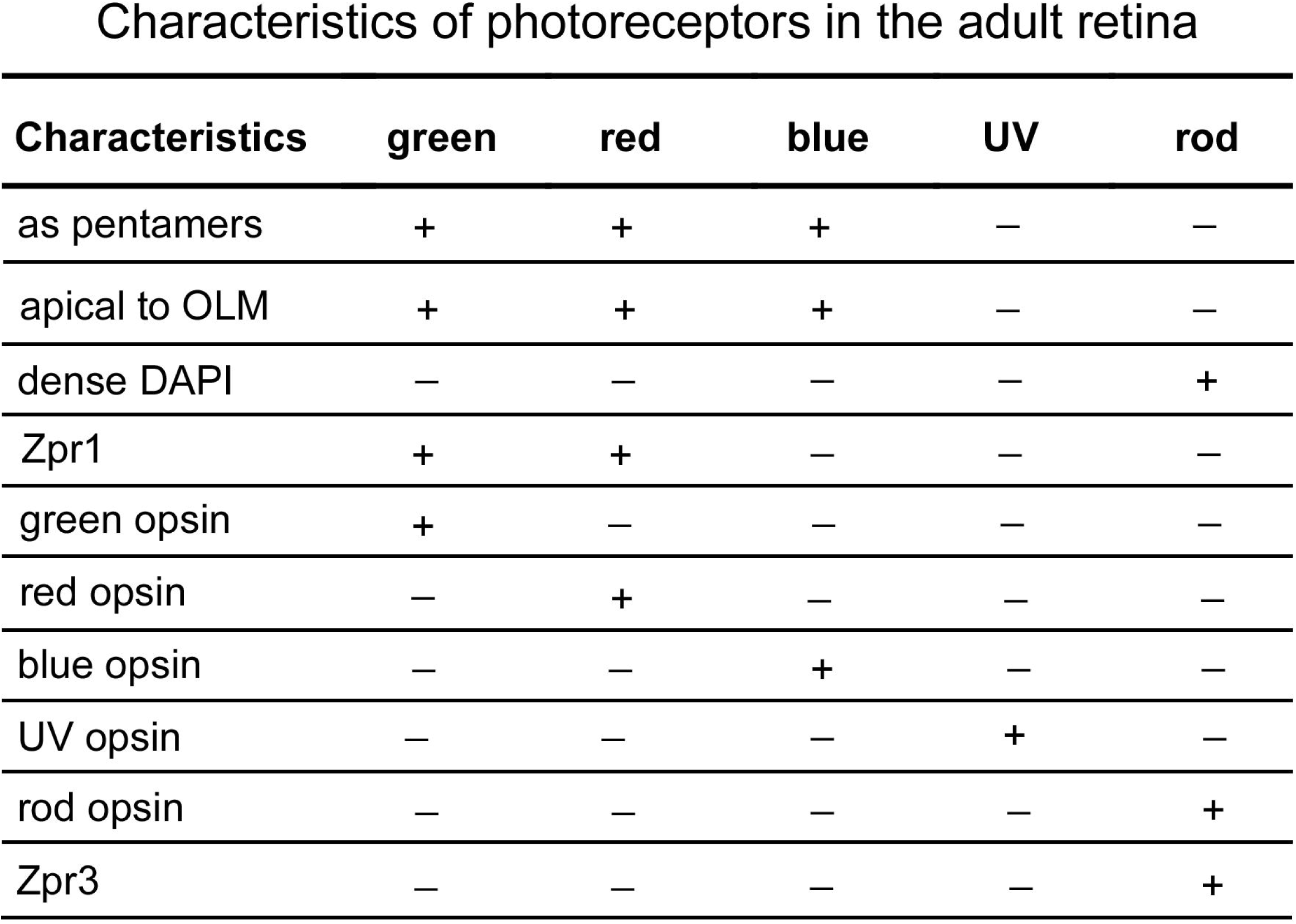
The table summarizes the morphological and immunoreactive characteristics of adult zebrafish photoreceptors. Plus signs stand for possession of the characteristics, and minus signs for nonpossession.

| Characteristics | green | red | blue | UV | rod |
| --- | --- | --- | --- | --- | --- |
| as pentamers | + | + | + | — | — |
| apical to OLM | + | + | + | — | — |
| dense DAPI | — | — | — | — | + |
| Zpr1 | + | + | — | — | — |
| green opsin | + | — | — | — | — |
| red opsin | — | + | — | — | — |
| blue opsin | — | — | + | — | — |
| UV opsin | — | — | — | + | — |
| rod opsin | — | — | — | — | + |
| Zpr3 | — | — | — | — | + |

**Table 2.**
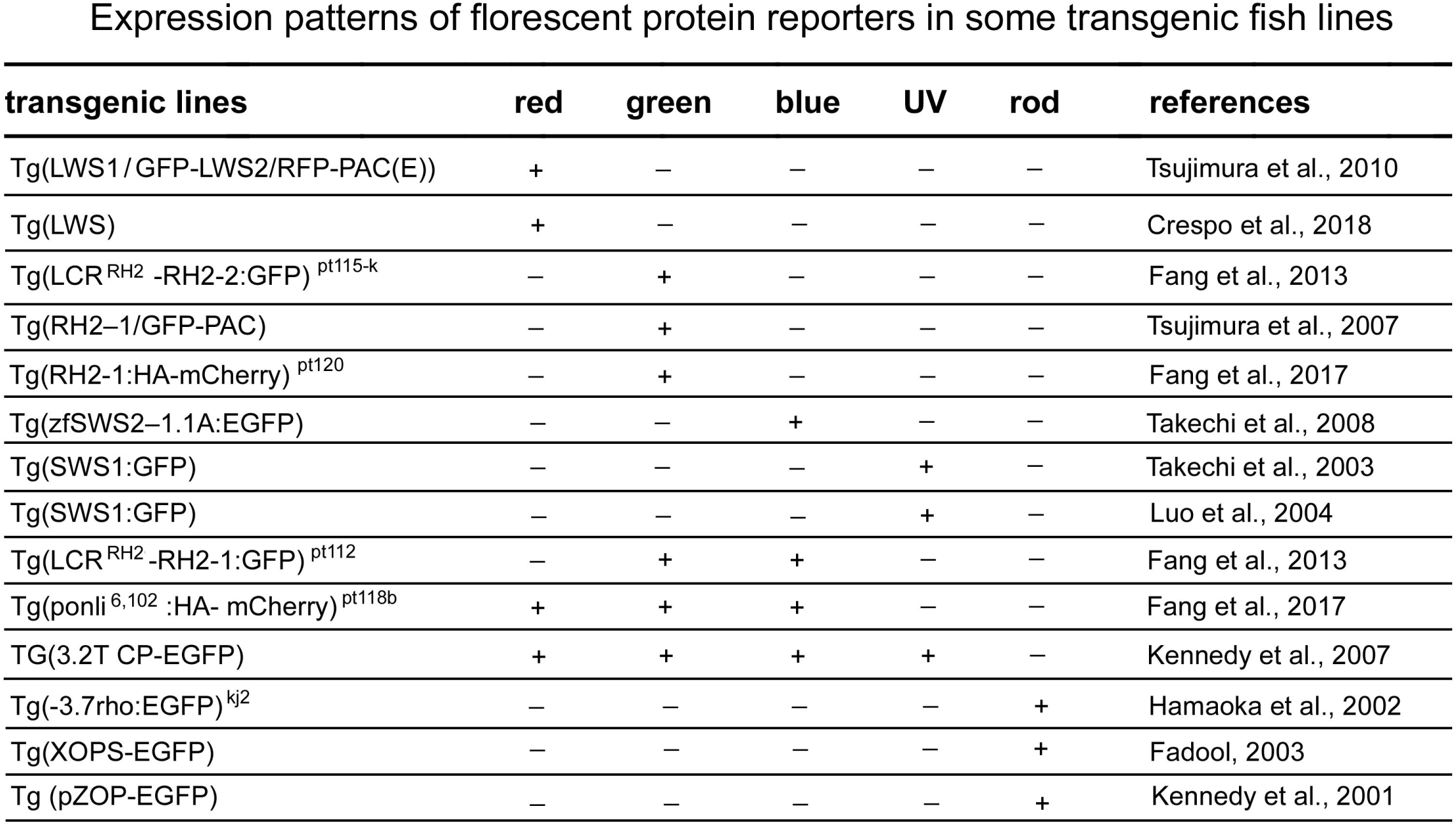
The table summarizes the expression patterns of fluorescent reporter proteins in the photoreceptors of some published transgenic zebrafish lines.

Expression patterns of florescent protein reporters in some transgenic fish lines
| transgenic lines | red | green | blue | UV | rod | references |
| --- | --- | --- | --- | --- | --- | --- |
| Tg(LWS1/GFP-LWS2/RFP-PAC(E)) | + | – | – | – | – | Tsujimura et al., 2010 |
| Tg(LWS) | + | – | – | – | – | Crespo et al., 2018 |
| Tg(LCR<sup>RH2</sup> -RH2-2:GFP)<sup>pt115-k</sup> | – | + | – | – | – | Fang et al., 2013 |
| Tg(RH2–1/GFP-PAC) | – | + | – | – | – | Tsujimura et al., 2007 |
| Tg(RH2-1:HA-mCherry)<sup>pt120</sup> | – | + | – | – | – | Fang et al., 2017 |
| Tg(zfSWS2–1.1A:EGFP) | – | – | + | – | – | Takechi et al., 2008 |
| Tg(SWS1:GFP) | – | – | – | + | – | Takechi et al., 2003 |
| Tg(SWS1:GFP) | – | – | – | + | – | Luo et al., 2004 |
| Tg(LCR<sup>RH2</sup> -RH2-1:GFP)<sup>pt112</sup> | – | + | + | – | – | Fang et al., 2013 |
| Tg(ponli<sup>6,102</sup> :HA- mCherry)<sup>pt118b</sup> | + | + | + | – | – | Fang et al., 2017 |
| TG(3.2T CP-EGFP) | + | + | + | + | – | Kennedy et al., 2007 |
| Tg(-3.7rho:EGFP)<sup>kj2</sup> | – | – | – | – | + | Hamaoka et al., 2002 |
| Tg(XOPS-EGFP) | – | – | – | – | + | Fadool, 2003 |
| Tg (pZOP-EGFP) | – | – | – | – | + | Kennedy et al., 2001 |

**Figure 4.**
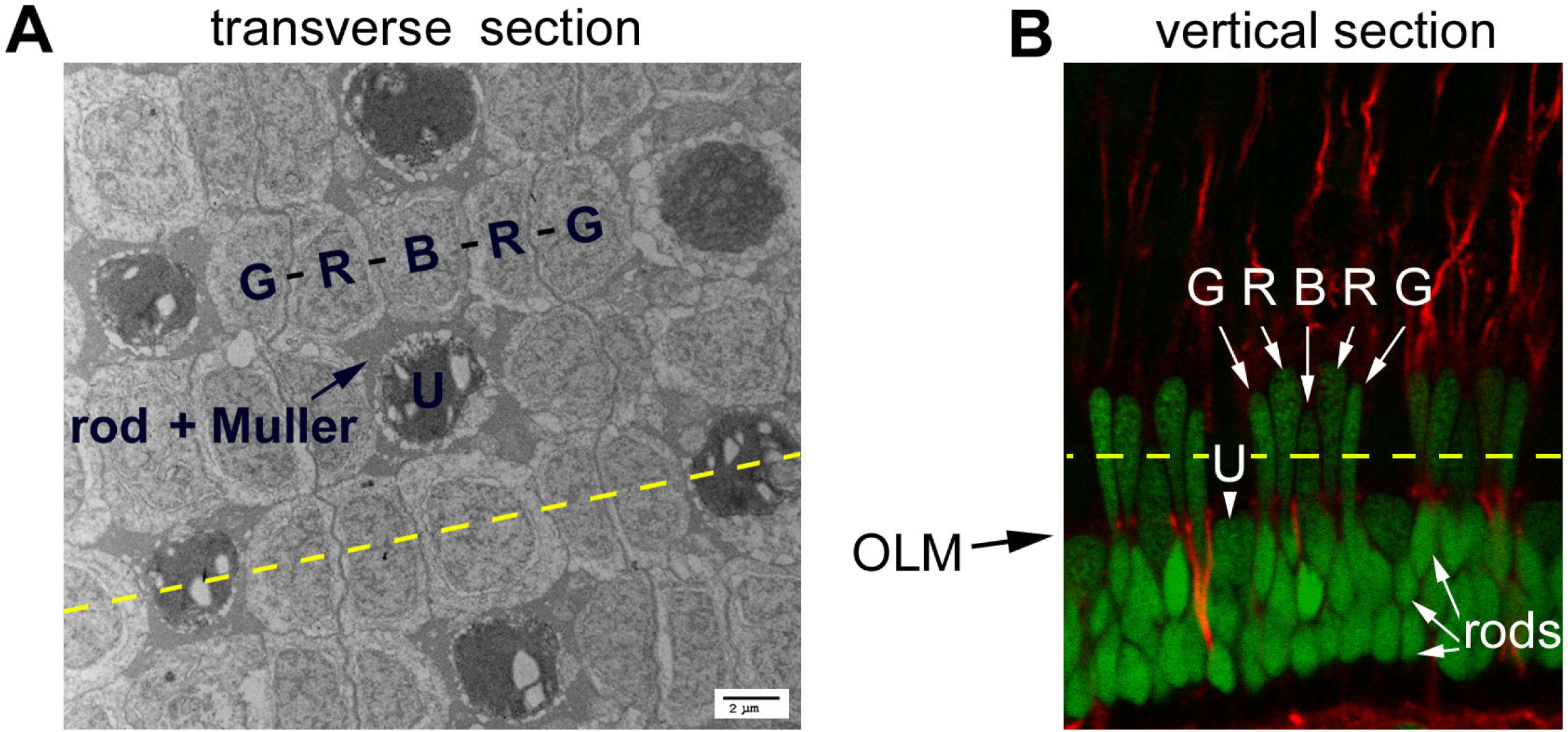
Morphological characteristics of five types of zebrafish photoreceptors. **A**. A TEM image of a transverse section of the photoreceptor cell layer of the adult zebrafish retina shows the mirror symmetric pentameric alignment of red, green, and blue (RGB) cones, indicated by G-R-B-R-G. U stands for UV cones, which display darkly-stained outer segments. An arrow indicates the clusters of the inner segments of rod photoreceptors and the apical processes of Müller glial cells around UV cones. The yellow dash line indicates the direction and location of vertical sectioning that would produce a retinal section as shown in panel B. **B**. A confocal immunohistochemical image of a vertical section of the adult zebrafish retina shows the morphologies and patterning of photoreceptor nuclei (by DAPI staining, green). Note that the elongated RGB cone nuclei locate apical to the outer limiting membrane (OLM), the round nuclei of UV cones (U) localize immediately basal to the OLM, and the strongly-stained rod nuclei locate away from OLM in the basal half of the outer nuclear layer. Red signals show the counterstaining of acetylated tubulin, some of which illustrate the OLM region. The yellow dashed line indicates the direction and position of transverse sectioning that would produce a retinal section as shown in panel A.

In the larval zebrafish retina, cones do not align in regular mosaics as in the adult retina (Allison et al., 2010). Thus, the planar arrangement of cones in the larval retina cannot be used as a criterion to distinguish photoreceptor types. However, the intensity of nuclear staining, immuno-reactivities, and comparison with existing transgenic zebrafish lines can still be utilized as criteria to distinguish photoreceptor types such as for the adult retina (Tables 2 and 3). In addition, the following morphological aspects of larval photoreceptors differ among photoreceptor types and can also be used as photoreceptor-type-identifying criteria: the positions of the nuclei, the position of the ellipsoid, and the cross-section sizes of inner segments at the OLM (Table 3; Fang et al., 2017).

**Table 3.**
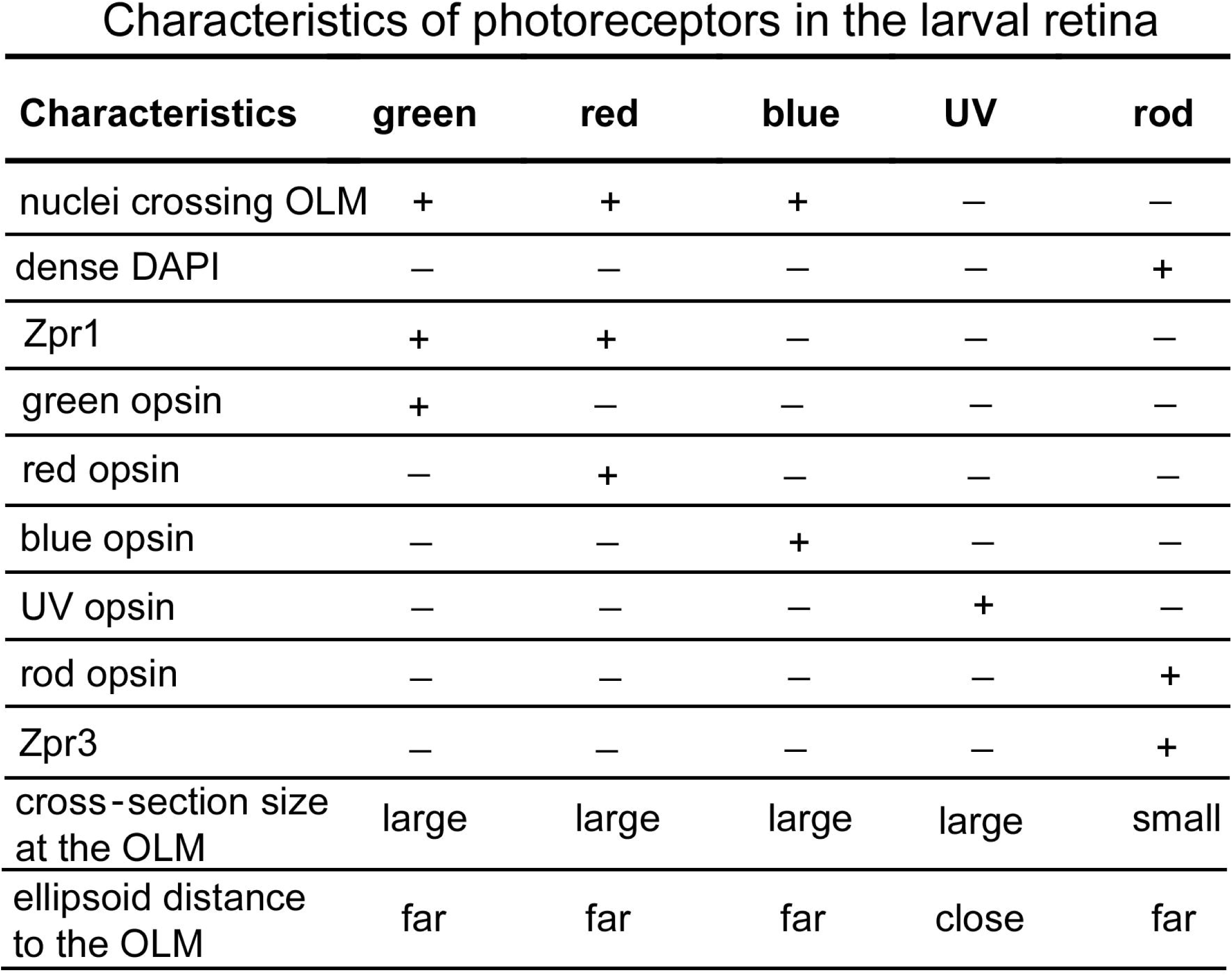
The table summarizes the morphological and immunoreactive characteristics of larval zebrafish photoreceptors at 9 dpf. Plus signs stands for possession of the characteristics, and minus signs for nonpossession.

Characteristics of photoreceptors in the larval retina
| Characteristics | green | red | blue | UV | rod |
| --- | --- | --- | --- | --- | --- |
| nuclei crossing OLM | + | + | + | — | — |
| dense DAPI | — | — | — | — | + |
| Zpr1 | + | + | — | — | — |
| green opsin | + | — | — | — | — |
| red opsin | — | + | — | — | — |
| blue opsin | — | — | + | — | — |
| UV opsin | — | — | — | + | — |
| rod opsin | — | — | — | — | + |
| Zpr3 | — | — | — | — | + |
| cross-section size at the OLM | large | large | large | large | small |
| ellipsoid distance to the OLM | far | far | far | close | far |

Although the above procedures are tailored for validating the transcriptional activity and specificity of the candidate core promoter and enhancer of a photoreceptor type-specific gene, by modifying the methods and criteria to assess GFP expression patterns, the procedure can also be used to validate the transcriptional activity of the CREs of other tissue-specific or cell type-specific genes.

### 4.3 Characterization of CREs

#### Rationale

Once the core promoter and enhancer regions of a gene are validated in Phase II, these CRE regions can be further characterized in Phase III. For example, by swapping CREs, deleting CREs, or mutating CRE motifs, one can further trim and narrow down the CRE regions, evaluate the contributions of individual CREs or CRE motifs to the transcriptional specificity of the gene, determine the functional conservation among orthologous CREs, or infer the grammatic code of the CREs. These analyses are explained as follows.

#### Procedures

##### 4.3.1 CRE swapping analysis

One way to dissect the functions of core promoters and enhancers is to analyze the expression patterns of reporter genes that are driven by the combination of the core promoter of one gene and the enhancer of another gene. The expression patterns driven by such chimeric combinations of CREs of two genes can then be compared with those by the CREs of a single gene to evaluate the roles of individual CREs in transcription.

For example, when the combination of the core promoter of the broadly-expressing *mpp5a* gene (also known as *nagie oko*; Wei and Malicki, 2002) and the enhancer of RGB cone-specific *mpp5b* gene drove RGB cone-specific transcription, we could infer that the *mpp5b* enhancer plays a critical role in RGB cone-specific transcription (Fang et al., 2017). Similarly, the RGB cone-specific transcription driven by the combination of zebrafish *mpp5b* core promoter and tilapia *mpp5b* enhancer suggests that the enhancers of *mpp5b* orthologs are conserved among teleost species (Fang et al., 2017).

##### 4.3.2 Deletion and mutation analyses

More drastic than the CRE swapping analysis, deletion analysis can be performed to remove or trim the core promoter and the enhancer of a transgenic GFP reporter construct. The resulting effects on transcriptional activity will shed light on the functions of the deleted regions. Moreover, the individual CRE motifs can be mutated to evaluate the roles of each motif in transcription. When mutating individual motifs, it is recommended that a motif be replaced with an unrelated sequence of the same length (namely, making substitution mutations rather than deletion mutations) because if the length of the CRE is altered by deletion, the spatial orientations of the neighboring binding sites for transcription factors and their intervals may be altered, potentially disrupting transcription factor interactions required for transcription, even if the motif sequence *per se* does not play a role in transcription.

When making a substitution mutation, a restriction enzyme site can be embedded in the substituting sequence. The inclusion of such an enzyme site allows for easy confirmation of the mutation by restriction digestion. Both deletion and substitution mutations can be achieved with the Q5 Site-directed Mutagenesis Kit (NEB, Cat #: E0554S).

## Acknowledgments

This work is supported by the National Institutes of Health [P30EY008098; EY025638; R21EY023665] as well as by the grants to the Department of Ophthalmology of the University of Pittsburgh from the Eye and Ear Foundation of Pittsburgh and Research to Prevent Blindness. The authors declare no competing financial interests. We thank Ms. Lynne Sunderman for proofreading the manuscript.

## References

Allison, W. T., Barthel, L. K., Skebo, K. M., Takechi, M., Kawamura, S., and Raymond, P. A. (2010). Ontogeny of cone photoreceptor mosaics in zebrafish. J Comp Neurol 518, 4182–4195.

Bailey, T. L., Boden, M., Buske, F. A., Frith, M., Grant, C. E., Clementi, L., Ren, J., Li, W. W., and Noble, W. S. (2009). MEME SUITE: tools for motif discovery and searching. Nucleic Acids Res 37, W202–208.

Bailey, T. L., and Elkan, C. (1994). Fitting a mixture model by expectation maximization to discover motifs in biopolymers. Proc Int Conf Intell Syst Mol Biol 2, 28–36.

Bailey, T. L., and Gribskov, M. (1998). Combining evidence using p-values: application to sequence homology searches. Bioinformatics 14, 48–54.

Beer, M. A., and Tavazoie, S. (2004). Predicting gene expression from sequence. Cell 117, 185–198.

Buchwalow, I., and Bocker, W. (2010). Immunohistochemistry: Basics and Methods, Springer).

Bulger, M., and Groudine, M. (1999). Looping versus linking: toward a model for long-distance gene activation. Genes Dev 13, 2465–2477.

Carninci, P., Sandelin, A., Lenhard, B., Katayama, S., Shimokawa, K., Ponjavic, J., Semple, C. A., Taylor, M. S., Engstrom, P. G., Frith, M. C., et al. (2006). Genome-wide analysis of mammalian promoter architecture and evolution. Nat Genet 38, 626–635.

Chalkley, G. E., and Verrijzer, C. P. (1999). DNA binding site selection by RNA polymerase II TAFs: a TAF(II)250-TAF(II)150 complex recognizes the initiator. Embo J 18, 4835–4845.

Chong, J. A., Tapia-Ramirez, J., Kim, S., Toledo-Aral, J. J., Zheng, Y., Boutros, M. C., Altshuller, Y. M., Frohman, M. A., Kraner, S. D., and Mandel, G. (1995). REST: a mammalian silencer protein that restricts sodium channel gene expression to neurons. Cell 80, 949–957.

Crespo, C., Soroldoni, D., and Knust, E. (2018). A novel transgenic zebrafish line for red opsin expression in outer segments of photoreceptor cells. Dev Dyn.

Das, M. K., and Dai, H. K. (2007). A survey of DNA motif finding algorithms. BMC Bioinformatics 8 Suppl 7, S21.

Elgar, G. (2009). Pan-vertebrate conserved non-coding sequences associated with developmental regulation. Brief Funct Genomic Proteomic 8, 256–265.

Fang, W., Bonaffini, S., Zou, J., Wang, X., Zhang, C., Tsujimura, T., Kawamura, S., and Wei, X. (2013). Characterization of transgenic zebrafish lines that express GFP in the retina, pineal gland, olfactory bulb, hatching gland, and optic tectum. Gene Expr Patterns 13, 150–159.

Fang, W., Guo, C., and Wei, X. (2017). rainbow enhancers regulate restrictive transcription in teleost green, red, and blue cones. J Neurosci.

Goodrich, J. A., and Tjian, R. (2010). Unexpected roles for core promoter recognition factors in cell-type-specific transcription and gene regulation. Nat Rev Genet 11, 549–558.

Grant, C. E., Bailey, T. L., and Noble, W. S. (2011). FIMO: scanning for occurrences of a given motif. Bioinformatics 27, 1017–1018.

Green, M., and Sambrook, J. (2012). Molecular Cloning: A Laboratory Manual, Forth edition edn).

Haberle, V., and Stark, A. (2018). Eukaryotic core promoters and the functional basis of transcription initiation. Nat Rev Mol Cell Biol.

Hardison, R. C. (2000). Conserved noncoding sequences are reliable guides to regulatory elements. Trends Genet 16, 369–372.

Hardison, R. C., and Taylor, J. (2012). Genomic approaches towards finding cis-regulatory modules in animals. Nat Rev Genet 13, 469–483.

Herold, M., Bartkuhn, M., and Renkawitz, R. (2012). CTCF: insights into insulator function during development. Development 139, 1045–1057.

Hughes, T. (2011). A handbook of transcripion factors, Springer).

Juven-Gershon, T., and Kadonaga, J. T. (2010). Regulation of gene expression via the core promoter and the basal transcriptional machinery. Dev Biol 339, 225–229.

Kennedy, B. N., Alvarez, Y., Brockerhoff, S. E., Stearns, G. W., Sapetto-Rebow, B., Taylor, M. R., and Hurley, J. B. (2007). Identification of a zebrafish cone photoreceptor-specific promoter and genetic rescue of achromatopsia in the nof mutant. Invest Ophthalmol Vis Sci 48, 522–529.

Kennedy, B. N., Vihtelic, T. S., Checkley, L., Vaughan, K. T., and Hyde, D. R. (2001). Isolation of a zebrafish rod opsin promoter to generate a transgenic zebrafish line expressing enhanced green fluorescent protein in rod photoreceptors. J Biol Chem 276, 14037–14043.

Kent, W. J. (2002a). BLAT--the BLAST-like alignment tool. Genome Res 12, 656-664.

Kent, W. J., Sugnet, C. W., Furey, T. S., Roskin, K. M., Pringle, T. H., Zahler, A. M., and Haussler, D. (2002b). The human genome browser at UCSC. Genome Res 12, 996–1006.

Krebs, J., Goldstein, E., and Kilpatrick, S. (2017). Levin’s Gene xii).

Kulaeva, O. I., Nizovtseva, E. V., Polikanov, Y. S., Ulianov, S. V., and Studitsky, V. M. (2012). Distant activation of transcription: mechanisms of enhancer action. Mol Cell Biol 32, 4892–4897.

Latchman, D. (2008). Eukaryotic Transcription Factors, Fifth Edition edn (London, Academic Press).

Latchman, D. (2015). Gene control, Garland Science).

Lenhard, B., Sandelin, A., and Carninci, P. (2012). Metazoan promoters: emerging characteristics and insights into transcriptional regulation. Nat Rev Genet 13, 233–245.

Lodish, H., Berk, A., Zipursky, S. L., Matsudaira, P., Baltimore, D., and Darnell, J. (2000). Regulatory Sequences in Eukaryotic Protein-Coding Genes. In Molecular Cell Biology (New York, W.H. Freeman).

Long, H. K., Prescott, S. L., and Wysocka, J. (2016). Ever-Changing Landscapes: Transcriptional Enhancers in Development and Evolution. Cell 167, 1170–1187.

Luo, W., Williams, J., Smallwood, P. M., Touchman, J. W., Roman, L. M., and Nathans, J. (2004). Proximal and distal sequences control UV cone pigment gene expression in transgenic zebrafish. J Biol Chem 279, 19286–19293.

Middendorf, M., Kundaje, A., Wiggins, C., Freund, Y., and Leslie, C. (2004). Predicting genetic regulatory response using classification. Bioinformatics 20 Suppl 1, i232–240.

Nelson, A. C., and Wardle, F. C. (2013). Conserved non-coding elements and cis regulation: actions speak louder than words. Development 140, 1385–1395.

Nikolov, D. B., Hu, S. H., Lin, J., Gasch, A., Hoffmann, A., Horikoshi, M., Chua, N. H., Roeder, R. G., and Burley, S. K. (1992). Crystal structure of TFIID TATA-box binding protein. Nature 360, 40–46.

Panne, D., Maniatis, T., and Harrison, S. C. (2007). An atomic model of the interferon-beta enhanceosome. Cell 129, 1111–1123.

Pennacchio, L. A., and Rubin, E. M. (2001). Genomic strategies to identify mammalian regulatory sequences. Nat Rev Genet 2, 100–109.

Raymond, P., Barthel, L., and Curran, G. (1995). Developmental patterning of rod and cone photoreceptors in embryonic zebrafish. Journal of Comparative Neurology 359, 537–550.

Robinson, J., Schmitt, E. A., Harosi, F. I., Reece, R. J., and Dowling, J. E. (1993). Zebrafish ultraviolet visual pigment: absorption spectrum, sequence, and localization. Proc Natl Acad Sci U S A 90, 6009–6012.

Rosen, J. N., Sweeney, M. F., and Mably, J. D. (2009). Microinjection of zebrafish embryos to analyze gene function. J Vis Exp.

Rouault, H., Santolini, M., Schweisguth, F., and Hakim, V. (2014). Imogene: identification of motifs and cis-regulatory modules underlying gene co-regulation. Nucleic Acids Res 42, 6128–6145.

Takechi, M., Hamaoka, T., and Kawamura, S. (2003). Fluorescence visualization of ultraviolet-sensitive cone photoreceptor development in living zebrafish. FEBS Lett 553, 90–94.

Thermes, V., Grabher, C., Ristoratore, F., Bourrat, F., Choulika, A., Wittbrodt, J., and Joly, J. S. (2002). I-SceI meganuclease mediates highly efficient transgenesis in fish. Mech Dev 118, 91–98.

Tsujimura, T., Chinen, A., and Kawamura, S. (2007). Identification of a locus control region for quadruplicated green-sensitive opsin genes in zebrafish. Proc Natl Acad Sci U S A 104, 12813–12818.

Tsujimura, T., Hosoya, T., and Kawamura, S. (2010). A single enhancer regulating the differential expression of duplicated red-sensitive opsin genes in zebrafish. PLoS Genet 6, e1001245.

Vavouri, T., and Lehner, B. (2009). Conserved noncoding elements and the evolution of animal body plans. Bioessays 31, 727–735.

Vihtelic, T. S., Doro, C. J., and Hyde, D. R. (1999). Cloning and characterization of six zebrafish photoreceptor opsin cDNAs and immunolocalization of their corresponding proteins. Vis Neurosci 16, 571–585.

Warner, J. B., Philippakis, A. A., Jaeger, S. A., He, F. S., Lin, J., and Bulyk, M. L. (2008). Systematic identification of mammalian regulatory motifs’ target genes and functions. Nat Methods 5, 347–353.

Wei, X., and Malicki, J. (2002). nagie oko, encoding a MAGUK-family protein, is essential for cellular patterning of the retina. Nature Genetics 31, 150–157.

Wittkopp, P. J., and Kalay, G. (2011). Cis-regulatory elements: molecular mechanisms and evolutionary processes underlying divergence. Nat Rev Genet 13, 59–69.

Yanez-Cuna, J. O., Kvon, E. Z., and Stark, A. (2013). Deciphering the transcriptional cis-regulatory code. Trends Genet 29, 11–22.

Yuan, S., and Sun, Z. (2009). Microinjection of mRNA and morpholino antisense oligonucleotides in zebrafish embryos. J Vis Exp.

Zou, J., Yang, X., and Wei, X. (2010). Restricted localization of ponli, a novel zebrafish MAGUK-family protein, to the inner segment interface areas between green, red, and blue cones. Invest Ophthalmol Vis Sci 51, 1738–1746.

